# Pharmacological and Physiological Characteristics of Synaptic Transmissions from the Medial Prefrontal Cortex onto Noradrenergic Neurons and Their Presynaptic Neurons in the Mouse Locus Coeruleus

**DOI:** 10.1101/2025.11.07.687181

**Authors:** Pin-Huan Lai, Wei-Chen Hung, Ming-Yuan Min, Hsiu-Wen Yang

**Affiliations:** Department of Life Science, College of Life Science, National Taiwan University, Taipei 10617, Taiwan; The Center for Neurobiology and Cognitive Science, National Taiwan University, Taipei 10051, Taiwan; Departments of Biomedical Sciences, Chung-Shan Medical University, Taichung 40201, Taiwan; Departments of Medical Research, Chung-Shan Medical University, Taichung 40201, Taiwan

**Author notes:** Equal contributions (Ming-Yuan Min, Hsiu-Wen Yang).

## Abstract

The locus coeruleus (LC) is the primary source of norepinephrine in the brain and is known to modulate brain-wide arousal state. Recent evidence suggests that it also regulates immediate attentional responses by resetting related cortical networks to optimize behavioral outcomes. Cortical regions of high cognitive function, such as the medial prefrontal cortex (mPFC), are theorized to directly influence LC output for the purpose of behavioral regulation. However, the available evidence is insufficient to provide a comprehensive understanding of the underlying mechanisms and properties. To provide further comprehensive data on this issue, we combined *ex vivo* whole-cell recording with an optogenetic approach to study the synaptic transmission of mPFC inputs to LC neurons, including noradrenergic (NA) neurons and GABAergic neurons presynaptic to them (preLC neurons). Our findings indicate that the mPFC exhibits monosynaptic connections with both NA and GABAergic preLC neurons. These synaptic connections demonstrate cell-type-specific disparities in glutamate release properties. In comparison to those on GABAergic preLC neurons, the mPFC fibers synapsing on LC-NA exhibit a lower release probability (higher paired-pulse ratio) and demonstrate a presynaptic enhancement of glutamate release efficacy during behavior. The features of simultaneous connections onto LC-NA and GABAergic preLC neurons, which exhibit cell-type-specific differences in plastic function of the transmitter release, enable the mPFC to effectively multiplex information to the LC for the adaptive regulation of behavior.

## Introduction

The locus coeruleus (LC) is located in the dorsal pons and consists primarily of noradrenergic (NA) neurons (Swanson & Hartman, 1975). LC-NA neurons have diverse and extensive axonal projections throughout the brain and spinal cord (Aston-Jones & Waterhouse, 2016; Schwarz et al., 2015; Breton-Provencher & Sur, 2019) and are considered the sole source of norepinephrine in the forebrain. Due to their widespread projections and tonic firing activity, which varies to regulate arousal (Hobson et al., 1975; Aston-Jones & Bloom, 1981; Takahashi et al., 2010; Berridge et al., 1993, 2018; Carter et al., 2010), LC-NA neurons are believed to play an important role in regulating vigilance and the sleep-wake cycle. However, recent evidence has revealed a more intricate and specific role for these neurons in regulating behavior (Aston-Jones & Cohen, 2005; Bouret & Sara, 2005; Poe et al., 2020; Grimm et al., 2024). In animals performing complex tasks, LC-NA neurons fire tonically at a moderate rate of about 3–5 Hz, which is necessary for maintaining cognitive arousal (Aston-Jones et al., 1997, 1998). These neurons can also respond with phasic activation to salient environmental stimuli, such as goal-related stimuli during a task. The phasic LC activation and the subsequent release of norepinephrine facilitates neuronal network processing and adaptive behavioral responses (Bouret & Sara, 2004, 2005; Clayton et al., 2004; Aston-Jones & Cohen, 2005; Bouret & Richmond, 2015; Vazey et al., 2018; Grimm et al., 2024; Wilmot et al., 2024).

The specific role for the LC in regulating behavior would require the NA neurons to receive various inputs for computation to maintain an adequate tonic firing rate to sustain arousal and to discharge phasic activation to optimize behavioral responses to salient stimuli. Inputs from the medial prefrontal cortex (mPFC), including the anterior cingulate cortex (ACC) and the orbitofrontal cortex (OFC), are particularly important for the LC to perform these roles, as these areas are known for evaluating reward (ACC) or cost (OFC) of a stimulus (Schultz et al., 2000; O’Doherty et al., 2002; Ito et al., 2003; Roesch & Olson, 2004; Sugrue et al., 2004; Aston-Jones & Cohen, 2005). Despite their importance, there is currently only anatomical evidence of connections between the mPFC and LC-NA neurons (Arnsten & Goldman-Rakic, 1984; Sesack et al., 1989; Aston-Jones & Cohen, 2005; Kuo et al., 2020). Regarding physiological data, the properties of synaptic transmission from the mPFC to the LC are controversial. Some studies indicate directly excitatory transmission (Jodo & Aston-Jones, 1997; Jodo et al., 1998), while others suggest that mPFC activation inhibits LC activity via local inhibitory neurons (Hervé-Minvielle & Sara, 1995; Breton-Provencher & Sur, 2019). A recent study by Barcomb et al. (2022) used brain slice preparation combined with optogenetics to directly examine the monosynaptic neurocircuit and report the glutamatergic transmission properties of this pathway. However, the exact manner in which the mPFC drives LC-NA neuron activation remains unknown.

In addition to NA neurons, a group of GABAergic neurons presynaptic to NA neurons has been identified (Aston-Jones et al., 2004; Jin et al., 2016; Breton-Provencher & Sur, 2019; Kuo et al., 2020). However, it is unclear whether the mPFC regulates LC activation via direct synaptic connections with these GABAergic neurons. In this study, we examined the functional connections between the OFC and NA neurons, as well as between the OFC and the GABAergic neurons presynaptic to the NA neurons in the LC. We also investigated how stimulating the mPFC with a paradigm that mimics cortical neuron activity in physiological conditions affects these two types of neurons.

## Material and Methods

### Animals

All animal experiments were approved by the Institutional Animal Care and Use Committees at National Taiwan University and Chung Shan Medical University. Every effort was made to minimize the number of animals used. Wild-type (WT, C57BL/ 6JNarl from National Laboratory Animal Center, Taipei, Taiwan) and GAD^GFP^ (Tsunekawa et al., 2005) mice of both sexes were used. The mice were housed and bred in a temperature-controlled vivarium under a 12-hour light/12-hour dark cycle. Food and water were available to the mice *ad libitum*.

### Stereotaxic Surgery for intracerebral viral vector infusion

Mice aged 4–8 weeks underwent stereotaxic surgery. After receiving an intraperitoneal (ip.) injection of ketamine (75 mg/kg) and xylazine hydrochloride (15 mg/kg) to induce anesthesia, the animals were secured in a stereotaxic apparatus. A small craniotomy was made over the left mPFC, and the dura was reflected. Adenosine associative virus (AAV1-syn-ChrimsonR-tdTomato; Addgene, Watertown, MA, USA; 59171) was infused into the left mPFC at three coordinates. The coordinates were as follows: 0.3 lateral, 2.34 anterior, and 1.8 doral, or 1.9 anterior and 0.9 dorsal to the Bregma. At each coordinate, 50–100 nL of the virus was infused. The infusion was performed using a glass pipette with a 60–80 μm tip opening connected to a 2.0 μL Hamilton syringe (Neuros Model 7002 KH, Hamilton, Reno, NV, USA). The virus was manually delivered at a rate of 10 nL/min.

### Preparation of Brain Slices

Brain slices containing the LC were prepared 4 weeks after the stereotaxic surgery described above. The mice were deeply anesthetized via ip. injection of urethane (1.3 g/kg), then perfused with 10 mL of ice-cold slicing solution through the cardiovascular system. The slicing solution contained the following (in mM): 92 N-methyl-D-glucose (NMDG), 2.5 KCl, 1.2 NaH□PO□, 10 MgSO□, 0.5 CaCl□, 20 HEPES, 30 NaHCO□, 25 glucose, 5 sodium ascorbate, and 3 sodium pyruvate. It was oxygenated with 95% O□ and 5% CO□ and had a pH of 7.3–7.4. The osmolality was adjusted to 315–320 mOsm using sucrose. The brain was rapidly dissected, and the brainstem was blocked and embedded in 2% agarose in the slicing solution. Coronal brain slices (350 µm thick) were cut using a vibratome (VT1000 S, Leica Biosystems, Nussloch, Germany), and the slice comprising the LC was collected and incubated in the slicing solution at room temperature for 30 minutes. The slice was cut into two halves, which were then transferred to a holding solution for an additional hour of incubation. This solution contained the same ingredients as the slicing solution, except 92 mM NaCl, 2 mM MgSO□, and 2 mM CaCl□ replaced 92 mM NMDG, 10 mM MgSO□, and 0.5 mM CaCl□, respectively. The remaining forebrain, which comprised the mPFC and was not used for slicing, was preserved in 4% paraformaldehyde in 0.1 M phosphate buffer (PB) for a later histological examination of the virus injection sites. After at least one hour of recovery, one of the halves was transferred to a recording chamber mounted on an upright microscope (BX51WI, Olympus Optical Co., Ltd., Tokyo, Japan) equipped with Nomarski and epifluorescence optics and an ORCA-R2 camera (Hamamatsu Photonics, Shizuoka, Japan). The slice was continuously perfused with artificial cerebrospinal fluid (ACSF) at a rate of 1.5–2 mL/min. The ACSF contained the following (in mM): 119 NaCl, 2.5 KCl, 1 NaH□PO□, 1.3 MgSO□, 2.5 CaCl□, 26.2 NaHCO□, and 11 glucose. It was oxygenated with 95% O□ and 5% CO□ (pH 7.3–7.4, osmolality 305–310 mOsm). Both half-slices were used for recording.

### Electrophysiology

The recording procedures and identification criteria for LC-NA neurons were the same as those previously described (Wang et al., 2015; Kuo et al., 2020). In brief, the recorded LC-NA neurons were located in the LC proper and displayed delayed firing responses to supra-threshold current pulses when their membrane potentials were held at approximately −100 to −70 mV. For recording of GABAergic neurons presynaptic to LC-NA neurons, brain slices prepared from GAD^GFP^ mice were used. Neurons that expressed GFP and were in the ventromedial part of the LC and the peri-LC were selected for recording (see also Kuo et al., 2023). Whole-cell recordings of both types of neurons were performed under visual guidance using glass pipettes pulled from borosilicate glass capillaries (GC150F-10, Warner Instruments, Hamden, CT, USA). The pipettes had tip resistances of 10–15 MΩ when filled with an internal solution containing the following (in mM): 131 potassium gluconate, 2 KCl, 10 HEPES, 2 EGTA, 8 NaCl, 2 ATP, and 0.3 GTP (pH 7.2–7.3, osmolality 306 mOsm). For current-clamp recordings, the bridge balance of membrane voltage in response to current pulse injection (one-second duration) was made. Data were accepted only if the resting membrane potential (Vm) of the recorded cell was at least −40 mV and action potentials could overshoot 0 mV. For voltage-clamp recordings, Vm was clamped at −70 mV, unless otherwise specified. No compensation was made for series resistance when sampling excitatory postsynaptic currents (EPSCs). However, it was monitored continuously throughout the recording. Data were excluded if the series resistance varied by more than 30% of the average value, typically less than 20 MΩ for LC-NA neuron recordings and less than 40 MΩ for GAD^GFP^ neuron recordings. In a series of experiments, the cell-attached configuration of the patch-clamp recording method was used to sample the spontaneous action potential activity of GAD^GFP^ neurons. This was done to avoid disturbing the spontaneous firing patterns (see also Kuo et al., 2023). Recording was performed using a pipette filled with ACSF in voltage-clamp mode (Vm = 0). All recordings were performed at room temperature (25 °C) using a Multiclamp 700B amplifier (Molecular Devices, San Jose, CA, USA) unless otherwise specified. Signals were low-pass filtered at 2 kHz, digitized at 10 kHz, and analyzed using a Micro 1401 interface with Signal3 and Spike2 software (Cambridge Electronic Design, Cambridge, UK).

### Optogenetics and analysis of EPSCs

*Ex vivo* optogenetic stimulation of the mPFC axonal terminals was performed by illuminating the brain slice with blue light pulses (545–580 nm, 1.2 mW/mm² intensity, and 2 ms duration) unless otherwise specified. The Micro1401, running Signal3, controlled an OptoLED Light Source (Cairn Research, Faversham, UK) to generate light pulses. These pulses were delivered through the epifluorescence light path and the 40× water-immersion objective of the microscope (LUMPLFLN-40xW, Olympus, Tokyo, Japan). Excitatory postsynaptic currents (EPSCs) were elicited by a single light pulse or paired pulses with an interpulse interval of 50 ms every 15 seconds. These are referred to as eEPSCs. This stimulation paradigm was used to characterize the kinetics of eEPSCs and for pharmacological experiments. The eEPSC kinetics parameters were measured from averaged traces consisting of at least 40 sweeps. Amplitude was measured by subtracting the mean baseline current level (the mean minus three times its standard deviation, calculated over a stable 10 ms period before the light pulse) from the eEPSC peak. Latency was measured from the light pulse onset to the eEPSC onset, which was defined as the time at which the differentiated current level exceeded the mean plus three times the standard deviation of a one-second baseline. Rise time was measured as the time difference between 20% and 80% of the eEPSC amplitude. Decay time was measured as the time difference between the eEPSC peak and 36.8% of the eEPSC amplitude (an exponential decay). The paired-pulse ratio (PPR) was measured as the ratio of the eEPSC amplitude elicited by the second pulse to that elicited by the first pulse. In addition to the standard stimulation paradigm, physiologically relevant stimulation (PRS) was used in some experiments. This consisted of a five-minute train of 2 ms light pulses with an average frequency of 2 Hz, arranged in a random sequence.

### Biocytin Histochemistry and Immunohistochemistry

To examine the AAV injection site, the forebrain containing the mPFC, which was preserved in paraformaldehyde during brain sectioning, was stored at 4°C overnight. The tissue was then incubated in 30% sucrose in 0.1 M phosphate buffer (PB) at 4 °C overnight for cryoprotection. After incubation, the brain was cut into 50-μm-thick coronal sections using a freezing microtome. The sections were mounted on slides using either RapiClear 1.47 (SUNJin Lab, Hsinchu, Taiwan) or Fluoroshield Mounting Medium with DAPI (Cat. # AB104139, Abcam, Cambridge, UK), then examined using a Zeiss LSM 780 confocal microscope system (Carl Zeiss, Oberkochen, Germany). For *post hoc* validation of recorded neurons and virus transfection, brain slices were fixed with 4% paraformaldehyde in PB at 4°C overnight immediately after recording. The slices were briefly rinsed with PB and then incubated in phosphate-buffered saline (PBS, 0.9% NaCl in PB) for 10 minutes. Next, the slices were incubated in 2% bovine serum albumin in PBST (0.3% Triton X-100 in PBS) for one hour at room temperature to block non-specific binding. Slices from wild-type mice that were used to record LC-NA neurons were incubated with mouse anti-tyrosine hydroxylase (TH) (diluted 1:300; catalog #MAB318, EMD Millipore Corp., USA) and rabbit anti-RFP (diluted 1:2000; catalog #600401379, Rockland, PA, USA) in PBST at 4 °C overnight. After primary antibody incubation, the slices were incubated for two hours with a secondary antibody cocktail containing goat anti-mouse IgG conjugated to Alexa Fluor® 488 (1:200, catalog #115-545-003, Jackson ImmunoResearch, West Grove, PA), donkey anti-rabbit IgG conjugated to Alexa Fluor™ Plus 594 (1:600, catalog #A32754, Thermo Fisher Scientific, MA), and streptavidin-conjugated DyLight 405 (1:100, catalog #016-470-084, Jackson ImmunoResearch, West Grove, PA). Between each incubation step, the slices were washed with PBS at least three times for five minutes each. All incubations were performed at room temperature unless otherwise specified. For slices prepared from GAD^GFP^ mice that were used to record GABAergic pre-LC neurons, the aforementioned procedures were repeated, except the primary antibody cocktail was replaced with a cocktail of chicken anti-GFP (1:2,000 dilution, Cat. #ab13970, Abcam, Cambridge, UK) and rabbit anti-TH (1:1,000 dilution, Cat. #AB152, EMD Millipore Corp., USA). The secondary antibody cocktail was also replaced with a cocktail of donkey anti-chicken IgY conjugated to Alexa Fluor® 488 (1:200, Cat. #703-545-155, Jackson ImmunoResearch, West Grove, PA, USA) and donkey anti-rabbit IgG conjugated to Alexa Fluor™ Plus 594 (1:600, Cat. #A32754, Thermo Fisher Scientific, MA), and streptavidin-conjugated DyLight 405 (1:100; catalog #016-470-084, Jackson ImmunoResearch, West Grove, PA). After staining, the slices were mounted on gelatin-coated slides using RapiClear 1.47 and examined using a Zeiss LSM 780 confocal microscope.

### Statistical analysis

All statistical analysis was conducted using Excel (Microsoft, USA) or online calculators (Social Science Statistics and Statistics Kingdom). Data are presented as means ± SEM. Parametric tests (Student’s paired t-test) were employed for data with normal distribution. Otherwise, non-parametric analysis (Wilcoxon matched-pairs signed rank test and Mann-Whitney test) was used. The Shapiro-Wilk test was used to assess normality of distributions. Statistical significance was indicated as ∗P < 0.05, ∗∗P < 0.01, ∗∗∗P < 0.001. Sample sizes are denoted as n/N, representing the number of cells/number of animals.

## Results

### Optogenetic Study of Synaptic Transmission from mPFC to LC-NA Neurons

To investigate the properties of synaptic transmission from the mPFC to LC-NA neurons, WT mice were injected with an adeno-associated virus (AAV) carrying the syn-ChrimsonR-tdTomato gene in the left mPFC (Figure 1A). *Post hoc* histochemical analysis revealed that opsin expression was primarily in the prelimbic area (PrL) and/or the medial OFC (mOFC) (Figure 1B). The regions of AAV infection largely overlap with those reported by Kuo et al. (2020), who found the densest projections to the LC and peri-LC area compared to other prefrontal regions. Consistent with their findings, we observed a similar pattern of axonal projections from the unilateral mPFC onto dorsal pontine regions, including both sides of the LC (Figure 1C). Additionally, we found the densest mPFC fibers expressing ChrimsonR in the peri-LC region medial to the LC proper; the fiber density was sparse in other peri-LC regions (see red puncta in Figure 1C).

**Figure 1:**
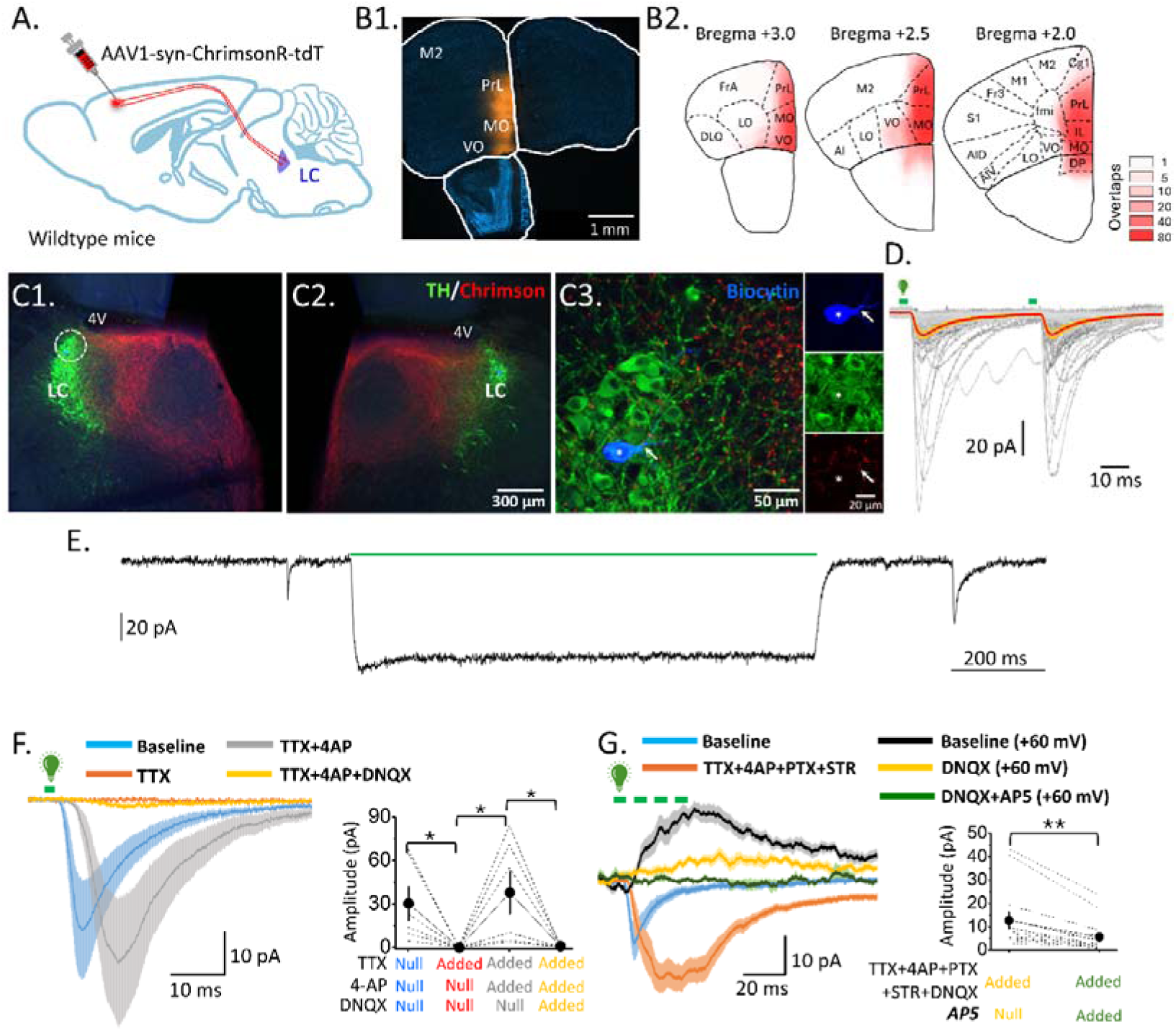
Optogenetic Study of Synaptic Transmission from mPFC to LC-NA Neurons. ***A.*** The schematic illustrates the AAV injection setup for the *ex vivo* simulation of the mPFC. ***B***. A representative case (B1) and a summed heatmap (B2) show that the AAV infusion sites are preliminary in the mPFC. *Aberration*: PrL: prelimbic cortex; MO: medial orbital cortex; VO: ventral orbital cortex. ***C.*** Fluorescent images from a representative experiment show the distribution of mPFC axonal fibers in the left (C1) and right (C2) brainstem slices comprising the LC at low power. The enlarged, circled regions of the dorsal left LC show a recorded LC-NA neuron (C3, asterisk), which was filled with biocytin (top right image) and exhibited TH-IR (middle right image). Note the surrounding axonal punctate (see arrows and the bottom-right inserted image). ***D.*** A representative experiment shows the elicitation of inward currents in an LC-NA neuron upon photostimulation with two pulses separated by a 50 ms interval. The green bars indicate the timing of the light pulses. The gray lines depict individual sweeps, the red line depicts the mean activity, and the yellow area depicts the standard error of the mean. Note that the response exhibits mild paired-pulse depression. ***E.*** A representative pharmacological experiment (left) and the summarized result (right) demonstrate that eEPSCs in LC-NA neurons, induced by optogenetic stimulation of mPFC fibers, are monosynaptic and AMPAR-mediated. ***F.*** A representative recording (left traces) and the summarized result (right plot) show that NMDARs also contribute to eEPSCs. In the representative recordings (*E* and *F*), the lines depict the mean activity of ten (E) and 12 (F) consecutive sweeps and the shaded areas depict the standard deviations of the means. In the summarized results, dashed lines show individual results, symbols show the mean, and capped lines show the standard error of the mean. Asterisks indicate a significant difference at the p < 0.05 (*) and p < 0.01 (**) levels.

A total of 233 neurons in the LC were identified as NA neurons because their online firing properties were consistent with established physiological criteria (Wang et al., 2015; Kuo et al., 2020). *Post hoc* analysis also revealed their immunoreactivity (IR) to the anti-TH antibody (Fig. 1C3). Stimulating mPFC fibers with a single light pulse elicited inward currents in 63% of LC-NA neurons (244 of 387 cells; n = 86 mice), and these currents had kinetics resembling a synaptic event (Figure 1D). An outward current was not elicited in all cases. Given that our recording conditions suggest that the inward currents are likely excitatory, we concluded that the elicited currents are eEPSCs. This conclusion was validated by a pharmacological study described below. Among the recorded LC-NA neurons, 5% expressed ChrimsonR, possibly due to trans-synaptic opsin transfer (Zingg et al., 2017). Consistent with this speculation, these neurons responded to stimulation with a one-second light pulse by exhibiting sustained but not transient inward currents mediated by the light-sensing ion channels (Figure 1E). The data were discarded, and all subsequent data were collected from LC-NA neurons that did not express ChrimsonR. Nevertheless, results from LC-NA neurons expressing ChrimsonR further support the argument that the quick, transient currents elicited in LC-NA neurons that do not express ChrimsonR are eEPSCs. Finally, the data presented below were collected from the LC ipsilateral and contralateral to the AAV injection site, though the AAV infusion was only administered to the left mPFC. No difference was found in the data collected from either LC site (Table 1). Furthermore, we found no difference in the data collected from animals of different sexes (Table 2).

**Table 1.** EPSCs evoked by stimulation of mPFC inputs in ipsilateral and contralateral LC-NA Neurons using optogenetics.

|  | **Ipsilateral** | **Contralateral** | **P value\*** |
| --- | --- | --- | --- |
| **Amplitude**<br>**(pA)** | 17.142 ± 2.870<br>(n=61 cells,<br>39 mice) | 13.590 ± 2.151<br>(n=68 cells,<br>47 mice) | 0.324 |
| **Latency**<br>**(ms)** | 5.236 ± 0.146<br>(n=61 cells,<br>39 mice) | 5.746 ± 0.179<br>(n=68 cells,<br>47 mice) | 0.070 |
| **Rise time**<br>**(ms)** | 2.090 ± 0.239<br>(n=61 cells,<br>39 mice) | 2.059 ± 0.195<br>(n=68 cells,<br>47 mice) | 0.795 |
| **Decay time**<br>**(ms)** | 10.992 ± 1.094<br>(n=61 cells,<br>39 mice) | 9.551 ± 0.517<br>(n=68 cells,<br>47 mice) | 0.689 |
| **Paired-pulse**<br>**ratio** | 1.061 ± 0.102<br>(n=44 cells,<br>27mice) | 0.883 ± 0.105<br>(n=35 cells,<br>26 mice) | 0.263 |

\*Mann-Whitney test.

**Table 2.** EPSCs evoked by stimulation of mPFC inputs in LC-NA Neurons using optogenetics in slices from male and female mice.

|  | Male | Female | P value* |  |
| --- | --- | --- | --- | --- |
| **Amplitude (pA)** | 16.199 ± 2.374<br>(n=89)<br>(n=89 cells,<br>37 mice) | 12.974 ± 2.109<br>(n=41 cells,<br>18 mice) | 0.719 | *Mann-Whitney test. |
| **Latency (ms)** | 5.574 ± 0.148<br>(n=89 cells,<br>37 mice) | 5.376 ± 0.194<br>(n=41 cells,<br>18 mice) | 0.548 |  |
| **Rise time (ms)** | 2.096 ± 0.191<br>(n=89 cells,<br>37 mice) | 2.000 ± 0.245<br>(n=41 cells,<br>18 mice) | 0.849 |  |
| **Decay time (ms)** | 10.685 ± 0.787<br>(n=89 cells,<br>37 mice) | 9.110 ± 0.688<br>(n=41 cells,<br>18 mice) | 0.352 |  |
| **Paired-pulse ratio** | 0.943 ± 0.082<br>(n=54 cells,<br>20 mice) | 1.066 ± 0.151<br>(n=25 cells,<br>12 mice) | 0.653 |  |

### EPSCs evoked in LC-NA neurons with optogenetics is monosynaptic and glutamatergic

An important advantage of studying synaptic transmission *ex vivo* combined with optogenetics is that the monosynaptic feature of the pathway can be tested directly (Petreanu et al., 2009). To this end, we examined the effects of tetrodotoxin (TTX, 1 μM), a voltage-gated Na□ channel blocker, and 4-aminopyridine (4-AP, 100 μM), a voltage-gated K□ channel blocker, on eEPSCs by optogenetics in LC-NA neurons (Figure 1F). The results showed that TTX completely abolished the eEPSCs (baseline vs. TTX: 30.541 ± 11.597 pA vs. 0.099 ± 1.356 pA; n/N = 6 cells/5 mice; p = 0.047, paired t-test). Subsequent application of 4-AP restored the eEPSCs (4-AP: 38.108 ± 14.785 pA; n = 6 cells/5 mice; p = 0.049 compared to TTX; paired t-test). Together, these results demonstrate that mPFC inputs have monosynaptic connections with LC-NA neurons.

We also tested the effects of two glutamate receptor antagonists: 6,7-dinitroquinoxaline-2,3-dione (DNQX, 10 μM), which blocks AMPA receptors (AMPARs); and (2R)-2-amino-5-phosphonopentanoic acid (AP5, 50 μM), which blocks NMDA receptors (NMDARs). With the Vm clamped at −70 mV, application of DNQX nearly eliminated the eEPSCs (DNQX: 1.026 ± 0.398 pA; n/N = 6 cells/5 mice; p = 0.050 compared to 4-AP; paired t-test; Figure 1F). To test the effect of AP5, we used a stronger photoactivation protocol (four light pulses at 100 Hz every 15 seconds). Additionally, to avoid interference from polysynaptic GABAergic and glycinergic activities, we applied TTX, 4-AP, picrotoxin (PTX, 100 μM), and strychnine (STR, 1 μM) into the bath medium. Finally, Mg²□ was removed from the bath medium, and the Vm was clamped at +60 mV to allow for more effective activation of NMDARs. Under these conditions, optogenetic activation of the mPFC fiber induced an outward current that was significantly reduced by AP5 (baseline: 12.671 ± 3.521 pA vs. AP5: 5.559 ± 1.850 pA; n/N = 14 cells/9 mice; p = 0.001; Wilcoxon matched-pairs signed rank test). This indicates a clear contribution of NMDA receptors (see Figure 1G). Together, these results confirm Barcomb et al.’s (2022) finding that EPSCs of mPFC inputs onto LC-NA neurons are mediated by glutamate acting at both AMPARs and NMDARs.

### Synaptic transmission from the mPFC to LC-NA neurons increases during behavior

During decision-making tasks, single-unit recordings from the preL cortex (part of the mPFC) show that neurons do not respond transiently or in an all-or-none manner to salient stimuli. Instead, they exhibit more persistent changes in activity at a rate of about 1–2 Hz over time (Bouret & Sara, 2004, 2005). To better understand how mPFC neuron activity under such physiological conditions regulates LC-NA neuron activity, we examined the effect on LC-NA neuron firing activity of a simulation paradigm of mPFC fibers that mimics preL unit activity during behavioral task. This paradigm is referred to as physiologically relevant stimulation (PRS). PRS consisted of a 5-minute train of 2-ms light pulses delivered at an average frequency of 2 Hz in a random sequence (Figure 2A). PRS increased synaptic transmission from the mPFC to LC-NA neurons compared to the resting condition (light pulses every 15 seconds as in pharmacological experiments; see Figure 1). This increase was associated with a reduction in transmission failure rate (Figure 2B1), defined as the proportion of light pulses failing to elicit eEPSCs. The transmission failure rate decreased from 0.302 ± 0.073 in the control condition to 0.113 ± 0.033 during PRS (n = 15 cells/11 mice; p = 0.0015; paired t-test; Figure 2C1). eEPSC potency, defined as the mean amplitude of successful eEPSCs, was also significantly enhanced by PRS (control: 43.890 ± 9.357 pA vs. PRS: 56.191 ± 9.820 pA; n/N = 15 cells/11 mice; p = 0.0413; paired-t test; Figure 2B2, 2C2). The failure rate and potency of eEPSCs returned to control levels after PRS (failure rate: 0.233 ± 0.057; n/N = 11 cells/9 mice; p = 0.001 compared to PRS; potency: 39.197 ± 7.479 pA; n/N = 11 cells/9 mice; p = 0.002 compared to PRS; paired t-test; Figure 2C1, 2C2).

**Figure 2.**
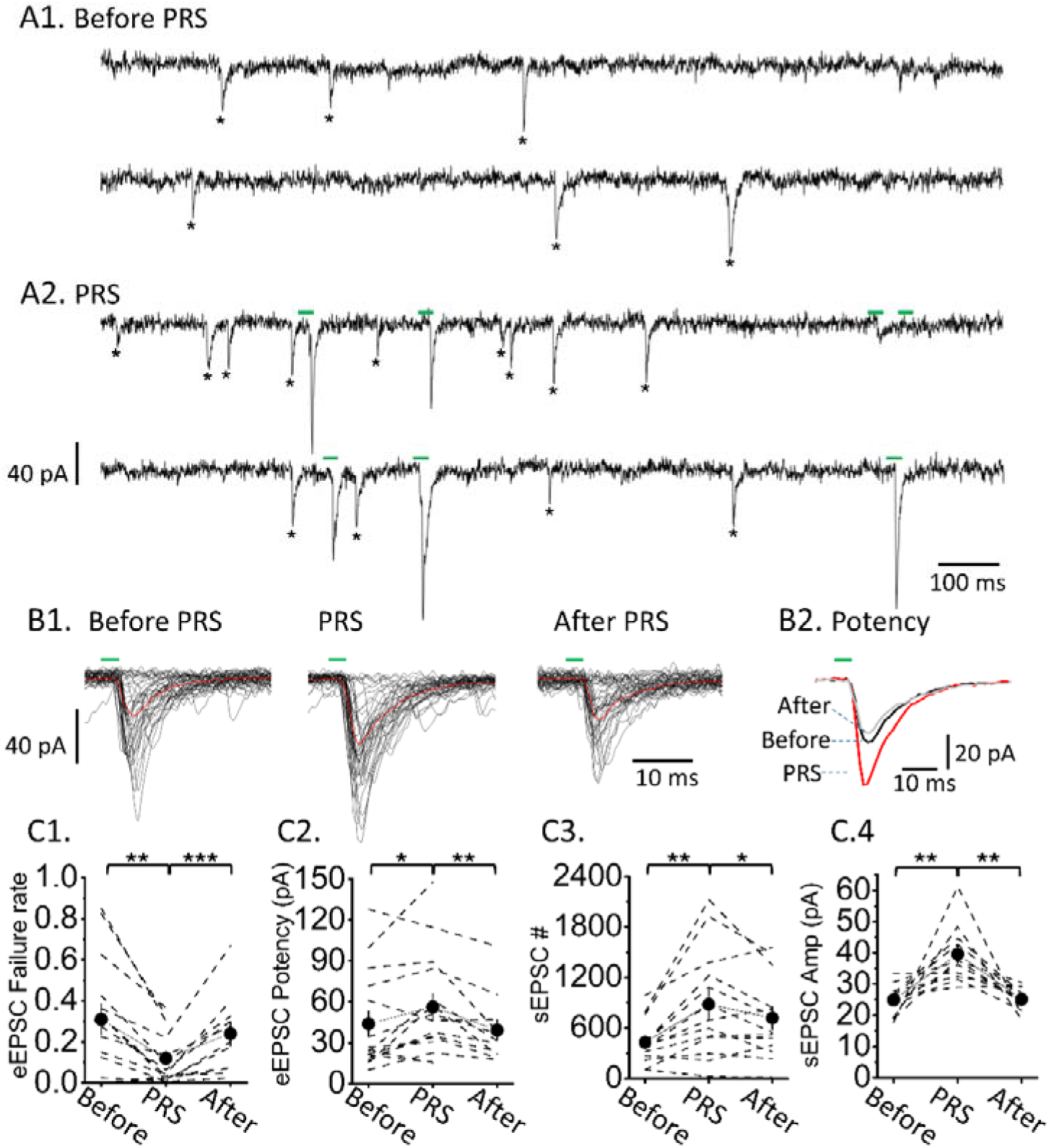
RPS increases synaptic transmission from the mPFC to LC-NA neurons. ***A.*** Recordings from a representative experiment show increased eEPSC and sEPSC (aEPSC) activity by RPS. The upper two traces show sEPSC activity before RPS (marked by an asterisk). The bottom two traces are recordings during RPS. The green bars mark the light pulses and the timed eEPSCs, and the asterisks mark the sEPSCs (aEPSCs). Note the random sequence of the light pulses. ***B.*** Superimpositions of 40 eEPSC sweeps in a representative experiment before RPS (elicited every 15 seconds; see the left traces in B1), during RPS (see the middle traces in B1), and 5 minutes after RPS (elicited every 15 seconds; see the right traces in B1) demonstrate successful and failure transmission. The red line shows the mean, and the black lines show individual sweeps. Excluding the sweeps of transmission failure shows the eEPSC potency of the three conditions (B2). The traces show the mean activity before, during, and after PRS. ***C.*** Plots show the summarized results of the effect of PRS on the failure rate (C1) and the potency of eEPSCs (C2), the number of sEPSCs (C3), and the mean amplitude of sEPSCs (C4). Dashed lines show individual results, symbols show the mean, and capped lines show the standard error of the mean. Asterisks indicate significant differences at the p < 0.05 (*) and p < 0.01 (**) levels.

In addition to eEPSCs, PRS increased spontaneous EPSCs (sEPSCs) during stimulation (Figures 2A and 2B). Analysis of 5-minute epochs before PRS revealed significant increases in both the number (control: 429.6 ± 74.0 vs. PRS: 879.6 ± 178.7, n = 13 cells/11 mice, p = 0.003; paired t-test; Figure 2C3) and mean amplitude of sEPSCs (control: 25.682 ± 1.439 pA vs. PRS: 37.426 ± 1.977 pA, n/N = 10 cells/8 mice, p = 0.0024; paired t-test; Figures 2C4). The number and mean amplitude of sEPSCs also returned to control levels after PRS (number: 453.5 ± 116.7; n/N = 13 cells/11 mice; p = 0.0266 compared to PRS; amplitude: 25.669 ± 1.141 pA; n/N = 10 cells/8 mice; p = 0.0017 compared to PRS; paired t-test; Figure 2C3-4), indicating that the increased sEPSC activity originated from mPFC stimulation, possibly due to asynchronous glutamate release. Together, these results show that PRS, which mimics physiological mPFC activity, enhances synaptic transmission from mPFC to LC-NA neurons, resulting in short-term potentiation by increasing presynaptic glutamate release.

Since LC-NA neurons exhibit spontaneous firing (Wang et al., 2015; Kuo et al., 2020), we next examined whether PRS of mPFC inputs influences the pattern of their spontaneous firing in current-clamp mode. PRS did not significantly change the mean firing rate of LC-NA neurons (control: 1.11 ± 0.30 Hz vs. PRS: 0.93 ± 0.22 Hz; n/N = 9 cells/5 mice; p = 0.2031; Wilcoxon matched-pairs signed rank test; Figures 3A–B). However, PRS clearly disrupted firing regularity, as reflected by a significant increase in the coefficient of variation of inter-spike intervals (ISI-CoV) (control: 0.33 ± 0.10 vs. PRS: 0.47 ± 0.14; n/N = 9 cells/5 mice; p = 0.0078; Wilcoxon matched-pairs signed rank test; Figure 3C). The ISI-CoV returned to baseline after PRS (0.34 ± 0.10; n/N = 9 cells/5 mice; p = 0.0039 compared to PRS; Wilcoxon matched-pairs signed rank test; Figure 3C). These results suggest that PRS of mPFC inputs could alter the firing pattern by increasing synaptic (membrane potential) noise in LC-NA neurons without significantly changing the overall firing frequency. Interestingly, each light pulse during PRS produced a brief depolarization (∼3 mV) followed by a prolonged hyper-polarization (up to 100 ms; Figure 3D).

**Figure 3.**
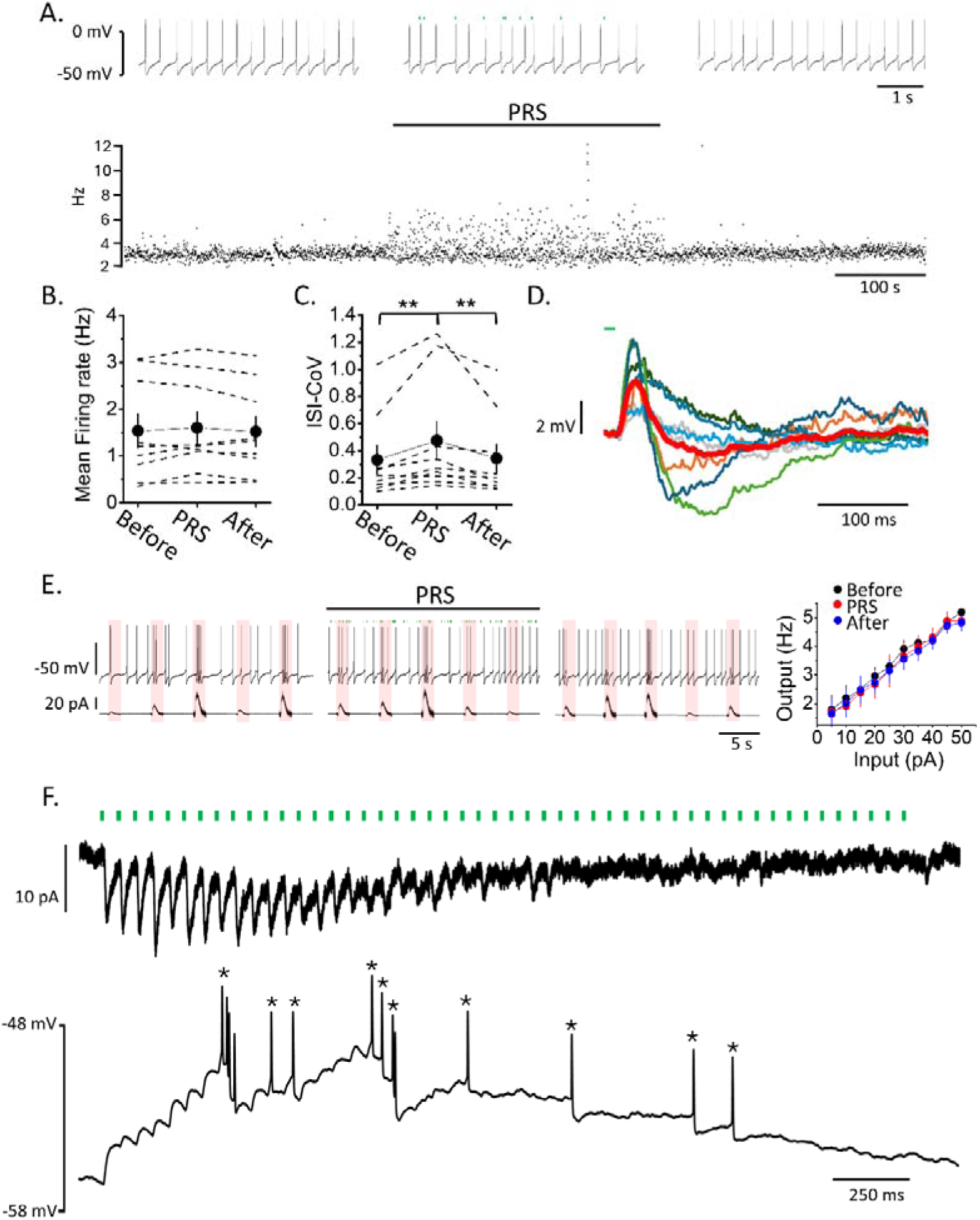
RPS disrupts the pattern, but not the mean firing rate, of spontaneous firing activity in LC-NA neurons. ***A.*** I-current recordings from a LC-NA neuron in a representative experiment demonstrate the impact of RPS on spontaneous AP activity. The top traces show the AP activity before (left), during (middle), and after (right) PRS. The bottom raster dot shows the instant firing frequency. The horizontal bar shows the duration of PRS. ***B, C*.** Plots show the summarized results of mean firing frequency (B) and ISI-CoV (C) analysis. The dashed lines show the individual results, the symbols show the mean, and the capped lines show the standard error of the mean. Asterisks indicate significant differences at the p < 0.05 (*) and p < 0.01 (**) levels. ***D*.** Superimposition of Vm responses triggered by light pulses during PRS in a representative experiment. The red trace shows the mean Vm response, and the other colored traces (7 traces) show the individual responses. ***E.*** Recordings from a representative experiment (left) and the summarized results (right) demonstrate the impact of PRS on the relationship between current injection intensity and phasic activation generation incidence. The recording traces in the top left panel show spontaneous AP before (left), during (middle), and after (right) PRS. The bottom left traces show the current injection templates of various intensities. The symbols and capped line in the right plot show the mean and standard error, respectively. ***F.*** Recordings from a representative experiment show the response of an LC-NA neuron to photostimulation at a high frequency of 20 Hz. The top trace shows a V-clamp recording, and the bottom trace shows an I-clamp recording of the same neuron. Green bars indicate photostimulation. Asterisks mark triggered APs by photostimulation. The responses shown here are averaged from 10 sweeps.

Increased synaptic noise has been shown to linearize the relationship between a neuron’s synaptic input and the generation of action potentials in thalamic neurons (Wolfart et al., 2005). Therefore, we examined whether PRS had a similar effect on LC-NA neurons. A notable aspect of recording from LC-NA neurons in whole-cell configuration is the ability to record large sEPSCs with high amplitude and long duration that can effectively drive LC phasic activation (Kuo et al., 2020). We used the averaged waveform of these large sEPSCs as a template for current injections (rise time: 108 ms; decay τ: 226 ms; duration: 760 ms; Figure 3E, bottom). A series of injections with varied intensities (5-50 pA) were delivered in a random sequence. Several series of injections were given, and the incidence of phasic activation generation by a given intensity was counted. We constructed the relationship between synaptic input and phasic activation output by plotting the current injection intensities against the incidence of phasic activation. Using this setup, we found that PRS did not change the slope of the input–output relationship (baseline: 0.075 ± 0.009 Hz/pA vs. PRS: 0.077 ± 0.005 Hz/pA; p = 0.6922; recovery: 0.073 ± 0.009 Hz/pA; p = 0.4254 compared to PRS; n/N = 6 cells/2 mice; paired t-test; Figure 3E). The results showing no change in firing rate (Figure 1A and 1B) or in the input-to-output relationship of LC-NA neurons by PRS of mPFC input motivate us to determine whether a higher mPFC stimulation frequency could change firing patterns of LC-NA neurons. First, we confirmed that well-timed eEPSCs could be triggered by a high-frequency (20 Hz) stimulation train, though rundown of the activity was also observed at the end of the stimulation train with voltage-clap recording. In the corresponding current-clamp recording, we indeed found that action potentials were triggered (Figure 3F). These results demonstrate that mPFC input can trigger LC-NA neuron firing activity when stimulated with an intensity beyond the physiological range.

### The synaptic transmission of mPFC onto LC-NA neurons is not influenced by Ptx + Stry

There is accumulating evidence demonstrating the existence of local GABAergic neurons that are presynaptic to LC-NA neurons (Aston-Jones et al., 2004; Jin et al., 2016; Breton-Provencher & Sur, 2019; Kuo et al., 2020; Luskin et al., 2025). They are referred to as GABAergic preLC neurons in this study. Previous studies have shown that GABAergic preLC neurons exert feedforward inhibition of LC-NA neurons (Kuo et al., 2020). They have also been shown to regulate arousal (Breton-Provencher & Sur, 2019; Luskin et al., 2025), pre-pulse inhibition in an LC-dependent manner (Kuo et al., 2020), and other physiological functions associated with the LC (Luskin et al., 2025). *In vivo* studies have suggested that mPFC activation suppresses LC-NA neuron activity through GABAergic preLC neurons (Hervé-Minvielle & Sara, 1995; Breton-Provencher & Sur, 2019). However, others *in vivo* (Jodo & Aston-Jones, 1997; Jodo et al., 1998) and *ex vivo* (Barcomb et al., 2022) studies have indicated that mPFC activation has a direct excitatory effect. To address this inconsistency, we investigated how blocking GABAergic and glycinergic transmissions affects eEPSCs in LC-NA neurons when the mPFC is optogenetically activated. To increase the likelihood of GABAergic preLC neuron activation, three light pulses at 20 Hz were used to photostimulate mPFC axonal fibers. We found that applying 100 μM picrotoxin (Ptx), a GABA-A receptor antagonist, and 1 μM strychnine (Stry), a glycine receptor antagonist, did not affect the mean amplitude of eEPSCs elicited by this stimulation paradigm (Figure 4A). The mean amplitudes of the eEPSCs were 15.2 ± 5.5 pA (first pulse), 30.3 ± 9.8 pA (second pulse), and 36.0 ± 8.4 pA (third pulse) prior to the application of PTX and STR. These values did not change significantly during drug application (first pulse: 18.3 ± 6.3 pA, p = 0.274; second pulse: 35.2 ± 8.9 pA, p =0.212; third pulse: 36.1 ± 8.0 pA, p = 0.986; n/N = 12 cells/5 mice; paired t-test; Figure 4A1, 4A2). Since the effect of the application of Ptx and Stry is likely bi-synaptic and could affect the decay of the eEPSCs more than their rise, we also measured the exponential decay time of the eEPSCs. There was no significant difference in the results (control vs. PTX + STR: first pulse: = 8.5 ± 1.0 ms vs. 9.7 ± 1.3 ms, p = 0.181; second pulse: 8.8 ± 0.9 ms vs. 8.5 ± 0.7 ms, p = 0.601; third pulse: 8.7 ± 1.1 ms vs. 9.1 ± 0.9 ms, p = 0.700; n/N = 12 cells/5 mice; Figure 4A3).

**Figure 4.**
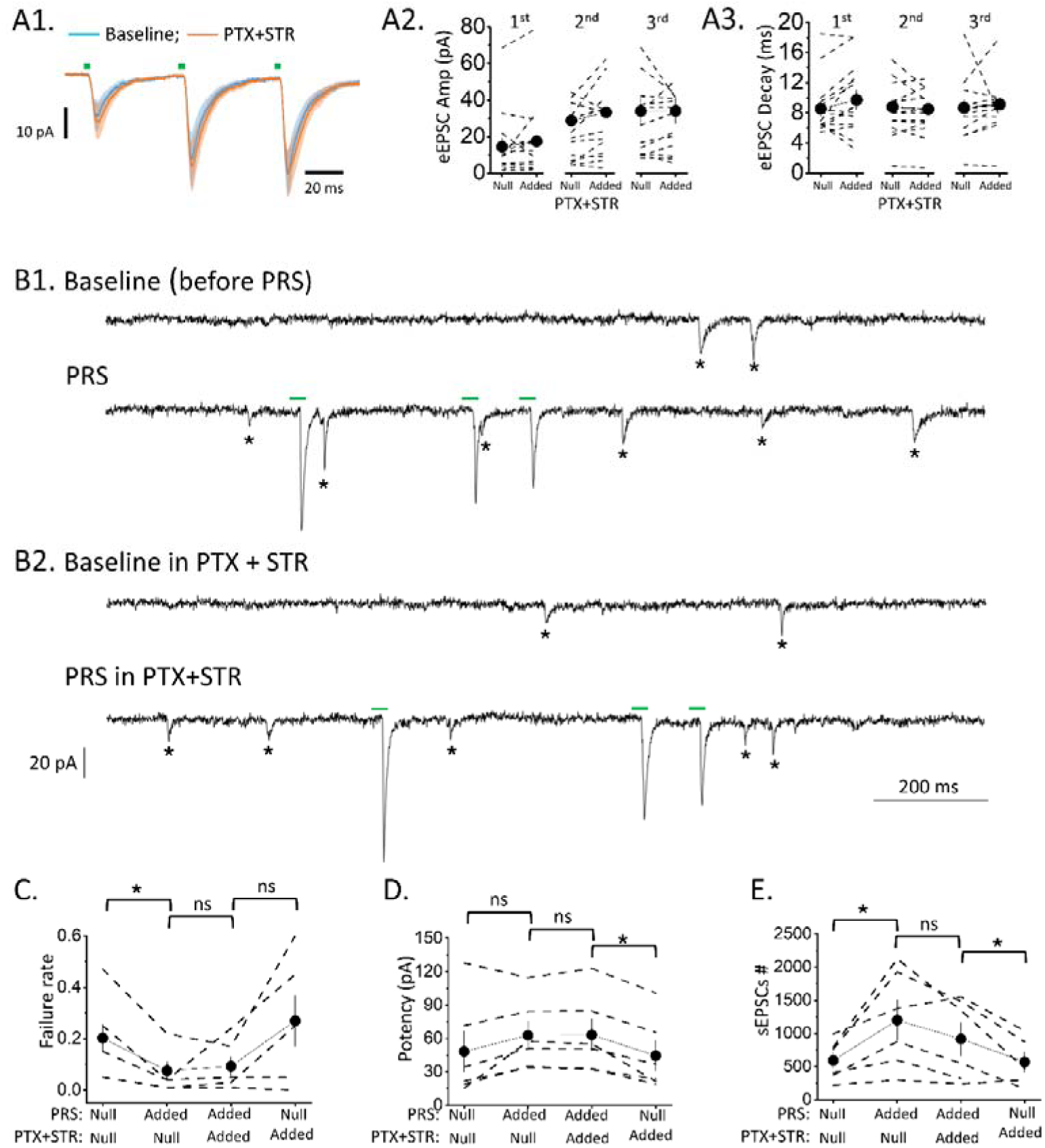
Application of Ptx + Stry does not influence the synaptic transmission of the mPFC onto LC-NE neurons. Representative recordings (A1) and summarized results of eEPSC amplitude (A2) and decay time (A3) analyses show that blocking the GABA-A receptor with picrotoxin (Ptx) and the glycine receptor with strychnine (Stry) had no effect on eEPSCs elicited in LC-NE neurons by optogenetic stimulation of mPFC fibers. In the representative recording (A1), the lines depict the mean activity of ten consecutive sweeps and the shaded areas depict the standard deviation of the means. In the summarized results (A2, A3), the dashed lines show the individual results, the symbols show the mean, and the capped lines show the standard error of the mean. ***B.*** Recordings from a representative experiment demonstrate increased eEPSC and sEPSC (aEPSC) activity by PRS before (B1) and during (B2) Ptx + Stry application. The upper and lower traces in panels B1 and B2 show recordings with and without RPS, respectively. Green bars mark light pulses and timed eEPSCs, and asterisks mark sEPSCs (aEPSCs). ***C-E***. The plots show the summarized results of the effect of PRS on the failure rate (C), the potency of eEPSCs (D), and the number of aEPSCs (E), with and without the application of Ptx + Stry. Dashed lines show individual results, symbols show the mean, and capped lines show the standard error of the mean. Asterisks indicate significant differences at the p < 0.05 (*) and p < 0.01 (**) levels. “ns” denotes no significance.

We also examined the effects of blocking GABAergic and glycinergic transmissions during PRS. Two PRS epochs were delivered, separated by a five-minute interval, with the drugs applied during halfway through the interval after the first PRS. PRS significantly reduced the failure rate (control vs. 1st PRS: 0.202 ± 0.052 vs. 0.076 ± 0.035, p = 0.019; n/N = 6 cells /5 mice; paired t-test; Figures 4B and C) and showed a trend of increasing EPSC potency (control vs. 1st PRS: 48.3 ± 18.0 pA vs. 62.7 ± 12.8 pA, p = 0.099; n/N = 6 cells/5 mice; paired t-test; Figures 4B and D). Further application of PTX and STR did not significantly alter the failure rate (2nd PRS: 0.093 ± 0.036, p = 0.671 compared to 1st PRS; n/N = 6 cells / 5 mice; paired t-test) and EPSC potency (2nd PRS: 63.1 ± 14.3 pA, p = 0.817 compared to 1st PRS; n/N = 6 cells/ 5 mice; paired t-test). The failure rate showed a trend of returning to control level after PRS under PTX and STR application (0.269 ± 0.098, p = 0.063 compared to 2nd PRS; n/N = 6 cells/5 mice; paired t-test). Meanwhile, the EPSC potency returned to control level after PRS (44.5 ± 13.3 pA, p = 0.003 compared to 2nd PRS; n/N = 6 cells/5 mice; paired t-test). Similarly, PRS increased sEPSC frequency (control vs. 1st PRS: 591.3 ± 122.2 vs. 1199.5 ± 300.2, p = 0.0354; n/N = 6 cells/6 mice; paired t-test; Figure 4E), and further PTX and STR application did not cause a significant effect (2nd PRS: 915.1± 249.8, p=0.086 compared to 1st PRS; n/N = 6 cells/ 6 mice; paired t-test). After PRS, sEPSC frequency returned to control level under PTX and STR application (568.8 ± 152.6, p = 0.040 compared to 2nd PRS; n/N = 5 cells/5 mice; paired t-test). Together, these results indicate that mPFC activation exerts a net excitatory effect on LC-NA neurons, and that GABAergic/glycinergic preLC neurons are not further activated by mPFC stimulation *ex vivo*, consistent with Barcomb et al. (2022).

### The synaptic transmission of mPFC to GABAergic pre-LC neurons

One possible explanation for the above findings is that the mPFC only has functional connections with LC-NA neurons and not with GABAergic preLC neurons. Alternatively, these connections may exist, but optogenetic stimulation of the mPFC alone may not be sufficient to change action potential pattern of GABAergic preLC neurons. To verify this, we investigated whether monosynaptic transmission exists from the mPFC to GABAergic pre-LC neurons (Figure 5A). Brain slices were prepared from GAD^GFP^ mice in which the promoter of GAD, an enzyme that synthesizes GABA, drives expression of GFP (Tsunekawa et al., 2005). We selected neurons that expressed GFP in the ventromedial part of the LC or peri-LC, a hotspot for GABAergic preLC neuron location (Breton-Provencher & Sur, 2019; Kuo et al., 2020; Luskin et al., 2025), for recording (Figure 5B). A total of 66 GABAergic preLC neurons were recorded. Compared to LC-NA neurons, in which 63% of recorded cells responded to mPFC activation, a significantly lower proportion of GABAergic preLC neurons responded (39%; 26 of 66 cells; see Figure 5C). To confirm that most of the recorded neurons were indeed preLC neurons, biocytin was injected into 32 of the 66 neurons during recording in a subset of experiments. The slices were stained with antibodies against GFP after the recording (Figure 5D). Although we selected neurons expressing GFP for recording, four of the injected neurons did not show post hoc GFP-IR. The loss of GFP-IR in these four cases could be due to cell content dialysis by the electrode solution during recording. Of the 28 neurons whose morphology was successfully retreated, we could identify axons with GFP-IR varicosities from 18 neurons that innervated the LC proper (Figure 5D). These results confirm that the majority of the recorded neurons are preLC neurons. All panels showing electrophysiological results in Figure 5 contain data from these 18 neurons.

**Figure 5.**
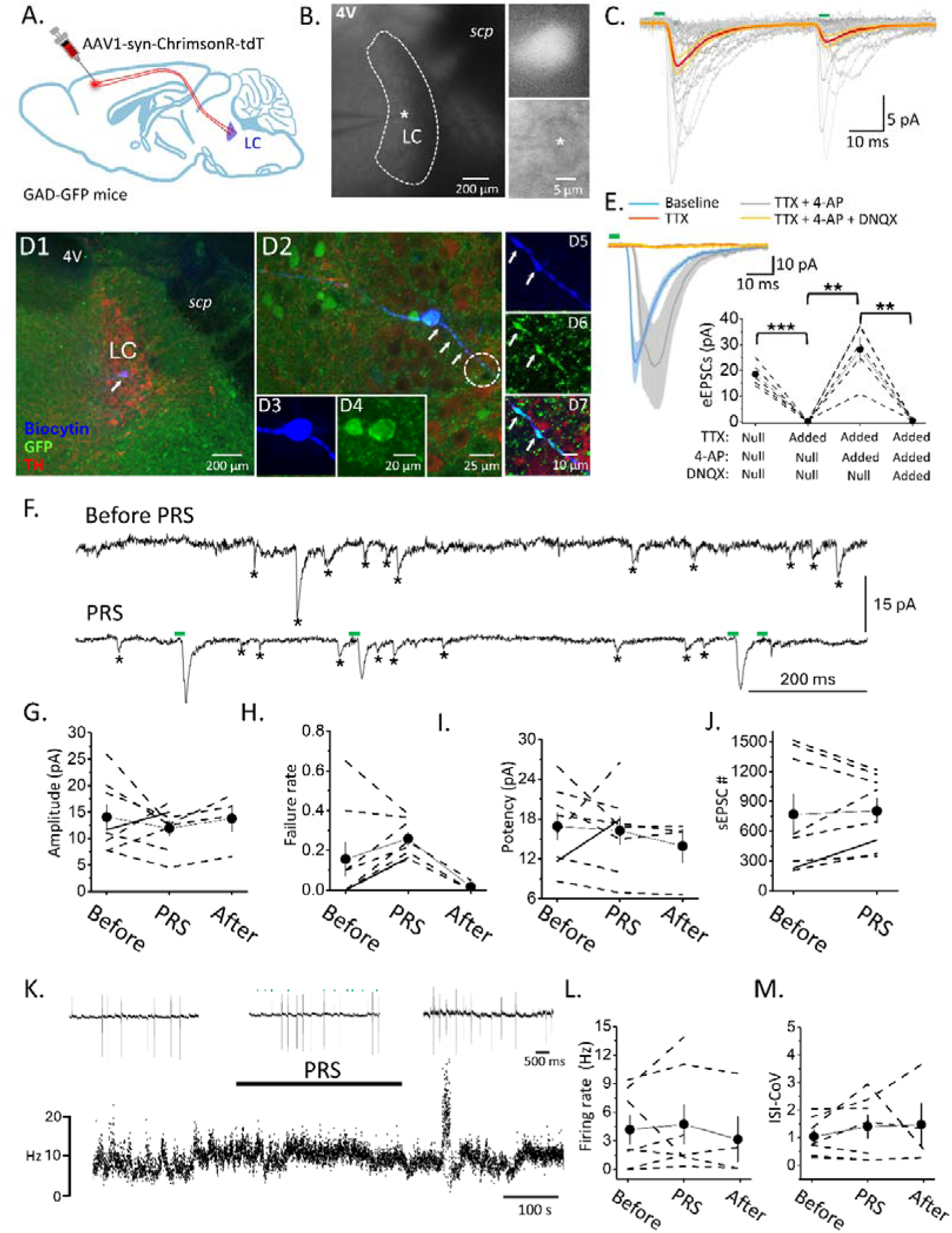
The synaptic transmission of mPFC to GABAergic pre-LC neurons. ***A.*** The schematic illustrates AAV injection setup for *ex vivo* recording of GABAergic pre-LC neurons and simulation of mPFC fibers. ***B.*** A representative online phase-contrast image shows the LC brain slice and the recording of a GABAergic pre-LC neuron. The asterisk marks the electrode tip, and the inserts on the right show an enlarged fluorescent image (top) and a phase-contrast image (bottom) of the electrode region. Note the recording from a neuron expressing GFP (inserts). ***C.*** A representative recording illustrates elicitation of inward currents from the same GFP expressing neuron (in B) upon photostimulation with two pulses separated by a 50 ms interval. The green bars indicate the timing of the light pulses. The gray lines depict individual sweeps, the red line depicts the mean activity, and the yellow area depicts the standard error of the mean. Note that the response exhibits paired-pulse depression. The green bar indicates the time of the light pulse. The gray lines show individual sweeps; the red line shows the mean activity; and the yellow line shows the standard deviation of the mean. ***D.*** Fluorescent images from the same experiment show *post hoc* validation of recorded GFP-expressing neuron that was filled with biocytin at low (D1) and high (D2) powers. Note that the neuron also shows GFP IR, which is more clearly seen in the further enlarged images (D3, D4). Also, note a branch of the axon (marked by arrows) in the LC. The images on the right show an enlargement of the circle region in D2, revealing the GFP-IR varicosities of the axonal branch (D5-D7). The GFP-IR varicosities make contact with TH-IR dendrites, indicating that the GFP-expressing neuron is a GABAergic pre-LC neuron. ***E.*** A representative pharmacological experiment (left) and its summarized results (right) demonstrate that eEPSCs elicited by optogenetic stimulation of mPFC fibers in GABAergic pre-LC-NA neurons are monosynaptic and AMPAR-mediated. ***F.*** Recordings from a representative experiment demonstrate eEPSC and sEPSC (aEPSC) recordings before (top) and during (bottom) PRS. Green bars mark light pulses and timed eEPSCs, and asterisks mark sEPSCs (aEPSCs). ***G–J***. The plots show the summarized results of the impact of PRS on the mean amplitude (G), failure rate (H), and potency of eEPSCs (I), and number of aEPSCs (J). ***K.*** Cell-attached recordings from a GABAergic pre-LC neuron in a representative experiment demonstrate the impact of PRS on spontaneous AP activity. The top traces show spike activity before (left), during (middle), and after (right) PRS. The bottom raster dot shows the instant firing frequency. The horizontal bar shows the duration of PRS. ***L. & M.*** Plots show summarized results of mean firing frequency (L) and ISI-CoV (M) analysis. Note that in all plots, the dashed lines show the individual results, the symbols show the mean, and the capped lines show the standard error of the mean. Asterisks indicate significant differences at the p < 0.05 (*) and p < 0.01 (**) levels.

Optogenetically evoked EPSCs in pre-LC neurons were abolished by TTX and restored by 4-AP (baseline vs. TTX: 18.5 ± 2.0 pA vs. 0.3 ± 0.1 pA; p = 0.001; 4-AP: 28.1 ± 4.9 pA; p = 0.005 compared to TTX; n/N = 5 cells/5 mice; paired t-test; Figure 5E), confirming monosynaptic connectivity. Further addition of DNQX eliminated the restored response (0.5 ± 2.0 pA; p = 0.0049 compared to 4-AP; n/N = 5 cells/5 mice; paired t-test; Figure 5E), indicating that transmission is glutamatergic. The EPSCs elicited in the LC-NA and preLC neurons by stimulation of mPFC inputs exhibit different kinetics (Table 3). Specifically, eEPSCs in preLC neurons exhibit significantly longer latencies and decay times. Additionally, preLC neurons have a significantly lower paired-pulse ratio than LC-NA neurons (see Figures 4A1 and 5E; Table 3). These results suggest that synaptic transmission from the mPFC to LC-NA and preLC neurons has different presynaptic properties; therefore, mPFC activation may exert differential effects on the two types of postsynaptic neurons in the LC. As expected, PRS of mPFC fibers did not significantly alter the mean amplitude of eEPSCs in preLC neurons (control vs. PRS: 14.0 ± 2.4 pA vs. 16.3 ± 2.1 pA; p = 0.4101; n/N = 8 cells/6 mice; paired t-test; Figures 5F & 5G), unlike in LC-NA neurons. This is consistent with the unaltered transmission failure rate (control vs. PRS: 0.19 ± 0.08 vs. 0.26 ± 0.03; n/N = 8 cells/6 mice; p = 0.369; paired-t test; Figure 5H) and potency of eEPSCs (control vs. PRS: 16.9 ± 2.1 pA vs. 16.3 ± 2.1 pA; p = 0.7794; n/N = 8 cells/6 mice; paired t-test; Figures 5I) in preLC neurons. Similarly, the number of sEPSCs did not show a significant difference during PRS (control vs. PRS: 769.4 ± 201.7 vs. 803.4 ± 127.9, p = 0.945; n/N = 8 cells/6 mice; Wilcoxon matched-pairs signed rank test; Figures 5F & 5I). Consistent with voltage-clamp recording results showing minor suppression or no effect on mPFC-to-preLC neuron synaptic transmission, PRS of mPFC fibers did not affect the mean firing rate (control: 4.18 ± 1.53 Hz vs. PRS: 4.74 ± 2.04 Hz; p = 0.469, n = 7 cells/6 mice; recovery: 3.16 ± 2.35 Hz; p = 0.375 compared to PRS; n/N = 4 cells/4 mice; Wilcoxon matched-pairs signed rank test; see Figure 5K & 5L) or ISI-CoV (control: 1.04 ± 0.26 vs. PRS: 1.40 ± 0.43; p = 0.21, n/N = 7 cells/6 mice; recovery: 1.48 ± 0.77; p = 0.723 compared to PRS; n/N = 4 cells/4 mice; paired t-test, see Figure 5K & 5M) in preLC neurons recorded with a cell-attached configuration. These findings are consistent with the results of the pharmacological experiment showing that the allocation of picrotoxin and strychnine had no effect on the transmission of synaptic signals from the mPFC to LC-NA neurons.

**Table 3.** Kinetics of eEPSCs elicited in LC-NA and GABAergic preLC neurons by optogenetic stimulation of mPFC fibers.

|  | **LC-NA** | **LC-GABA** | **P value\*** |
| --- | --- | --- | --- |
| **Amplitude**<br>**(pA)** | **15.182 ± 1.757**<br>(n=130 cells,<br>55 mice) | **8.086 ± 1.342**<br>(n=26 cells,<br>14 mice) | **0.097** |
| **Latency**<br>**(ms)** | **5.511 ± 0.118**<br>(n=130 cells,<br>55 mice) | **6.611 ± 0.374**<br>(n=26 cells,<br>14 mice) | **0.002** |
| **Rise time**<br>**(ms)** | **2.065 ± 0.151**<br>(n=130 cells,<br>55 mice) | **1.808 ± 0.304**<br>(n=26 cells,<br>14 mice) | **0.222** |
| **Decay time**<br>**(ms)** | **10.188 ± 0.583**<br>(n=130 cells,<br>55 mice) | **7.731 ± 0.795**<br>(n=26 cells,<br>14 mice) | **0.026** |
| **Paired-pulse ratio** | **0.982 ± 0.073**<br>(n=78 cells,<br>32 mice) | **0.577 ± 0.131**<br>(n=26 cells,<br>14 mice) | **<0.001** |

\* Matt-Whitney test.

## Discussion

In this study, we examined the properties of synaptic transmission from the mPFC to LC-NA neurons and GABAergic preLC neurons. Consistent with previous *ex vivo* studies (Barcomb et al., 2022), we demonstrate that LC-NA neurons receive monosynaptic transmission from the mPFC using a combination of optogenetic and pharmacological approaches (Petreanu et al., 2009). This protocol has been established as a method for identifying monosynaptic neuronal connections. Transmission from the mPFC to LC-NA neurons is mediated by glutamate acting at both AMPARs and NMDARs. In addition to stimulating the mPFC at a slow rate (every 15 seconds), a common stimulation paradigm for investigating the kinetics and pharmacological properties of synaptic transmission, we tested the effect of stimulating the mPFC under more physiologically relevant conditions. This stimulation paradigm emulates the neuronal activity of the mPFC in experimental animals engaged in a complex task (Bouret & Sara, 2004). Notably, in contrast to the *ex vivo* resting condition, we observed that synaptic transmission from the mPFC to LC-NA neurons appears to be more efficient through short-term presynaptic enhancement. This argument is based on two phenomena: a reduced transmission failure rate and an increase in sEPSC events. We posit that the augmented sEPSC events observed during RPS originate in the mPFC and are a consequence of asynchronous glutamate release. This proposition is substantiated by the RPS-dependent nature of the effect. Although an increase in EPSC potency and the mean amplitude of sEPSCs has been observed, postsynaptic modulation is considered an unlikely contributor to this process. This is because an increase in quantal content (a presynaptic factor) during RPS could contribute to the observed increase in eEPSC potency and sEPSC amplitude.

Despite the presynaptic enhancement, the mean firing rate of LC-NA neurons *ex vivo* remains unchanged by PRS of mPFC fibers. Conversely, the PRS disrupts the pattern of the spontaneous firing activity, as evidenced by increasing inter-spike interval variance. This observation is consistent with previous findings (de Oliveira et al., 2010; Wang et al., 2015; Kuo et al., 2020). The spontaneous action potential activity of LC-NA neurons *ex vivo* is irregular and sometimes exhibits burst activity when synaptic activity remains functional in the brain slices. Blocking both excitatory and inhibitory synaptic activity does not significantly change the mean firing rate of LC-NA neurons. However, it results in a very regular action potential pattern, akin to a pacemaker (Wang et al., 2015; Kuo et al., 2020). The interference of the action potential pattern may be due to the PRS generating substantial synaptic noise or membrane potential fluctuations in LC-NA neurons, consisting of a rapid depolarization followed by protracted hyperpolarization. Since the present study failed to demonstrate a substantial impact of Ptx plus Stry application on EPSC activity in LC-NA neurons elicited by a 3-pulse train in resting conditions (stimulated every 15 seconds) or by the PRS, it is improbable that the hyperpolarization component of the synaptic noise is mediated by GABAergic and/or glycinergic transmissions. Barcomb et al. (2022) corroborated the observation of no effect of the Ptx and Stry application.

LC-NA neurons exhibited a delay in firing action potentials in response to a depolarizing current pulse at supra-threshold intensity; the duration of this delay depended on the Vm (Wang et al., 2015; Kuo et al., 2020). This electrophysiological feature has been observed in other neurons that express A-type potassium current (I_K_) and is used as a signature of neuronal A-type I_K_ expression (Liss et al., 2001; Burdakov & Ashcroft, 2002; Burdakov et al., 2004). Indeed, the expression of A-type I_K_ in potine NA neurons has been confirmed (Min et al., 2010). A-type I_K_ in NA neurons is characterized by rapid activation and operation at subthreshold membrane voltages. Furthermore, it can be evoked by a voltage command that mimics EPSP activity in NA neurons. Consistent with these properties, blocking A-type I_K_ increases the likelihood that recorded NA neurons will generate action potential when glutamate is puffed onto them (Min et al., 2010). The expression of a substantial amount of A-type I_K_ in LC-NA neurons could provide a rationale for the observation of seldom timed action potentials, but not EPSCs, with the light pulses during the PRS. It is probable that the summation of excitatory synaptic potentials evoked by RPS of the mPFC was insufficient to trigger an action potential but sufficient to evoke A-type I_K_. This interrupted the action potential pattern generated by the intrinsic membrane properties of LC-NA neurons (de Oliveira et al., 2010), but it did not alter the mean firing rate. Although it can be triggered by high-frequency stimulation of the mPFC, the present results suggest that mPFC input alone cannot directly trigger action potentials within the physiological firing range. Given the massive mutual connections between the LC and various cortical regions, as well as the fact that numerous cortical regions are active during complex behavioral tests, it is meaningful for LC-NA neuron activation not to be driven by a single input but rather by the outcome of integrating functionally correlated inputs.

Another novel and significant finding of the present study is the monosynaptic contacts of mPFC fibers onto GABAergic preLC neurons. Despite their shared origin, the results of the present study suggest that the properties of glutamate release from mPFC axonal terminals exhibit cell-type-specific differences. Specifically, we compared LC-NA neurons and GABAergic pre-LC neurons. Transmission from the mPFC to LC-NA neurons exhibited significantly less paired-pulse depression than transmission from the mPFC to GABAergic preLC neurons (see Figures 1D and 5C), indicating a lower probability of glutamate release from the axonal terminals to LC-NA neurons than to GABAergic pre-LC neurons. Furthermore, mPFC-to-NA neuron transmission in the LC demonstrates presynaptic enhancement of release efficacy during the PRS. This phenomenon is absent in mPFC-to-GABAergic preLC neuron transmission (see Figures 2 and 5F–J). Target cell-specific differences in transmitter release at terminals from a single axon due to differential release probability and/or short-term plasticity (frequency-dependent facilitation) are not unique to this case, but rather a general feature of neuronal networks (Scanziani et al., 1998; Reyes et al., 1998; Tóth & McBain, 2000). Such a mechanism is hypothesized to enhance precise control of the excitation-inhibition balance, the timing of inhibition, and network gain control. Additionally, this mechanism facilitates temporal filtering because synapses that demonstrate frequency-dependent facilitation of transmitter release are adept at detecting burst activity. In contrast, those that exhibit depression are efficient for rapid signaling but less so for sustained input. After undergoing these processes, a presynaptic (mPFC) neuron can function as a frequency- or pattern-selective signal depending on its postsynaptic partners (LC-NA and GABAergic preLC neurons). Interestingly, GABAergic preLC neurons have been shown to coordinate phasic LC activation under cortical regulation (Breton-Provencher & Sur, 2019; Kuo et al., 2020; Luskin et al., 2025). However, we cannot confirm whether LC-NA and GABAergic preLC neurons receive axonal terminals from the same presynaptic neuron because data from these two types of neurons were not recorded simultaneously. Our injection site covered a large number of mPFC subregions. Watanabe et al. (2024) recently reported that the PrL and infralimbic regions of the mPFC project topographically to the LC and peri-LC regions and play differential roles in the extinction of emotional memories. Therefore, we cannot rule out the possibility that the difference in release probability and/or frequency-dependent facilitation between mPFC-to-LC-NA and mPFC-to-GABAergic preLC neuron transmissions is due to different types of presynaptic neurons in the mPFC.

Unlike LC-NA neurons, which respond at a rate of 63%, fewer recorded GABAergic pre-LC neurons respond to photostimulation of mPFC fibers. This may reflect the molecular and functional heterogeneity of this group of neurons and indicate that not all types of GABAergic pre-LC neurons receive mPFC inputs. Furthermore, unlike LC-NA neurons, PRS of the mPFC did not disrupt the spontaneous firing pattern of GABAergic preLC neurons by influencing the ISI-CoV. As our previous study showed, some GABAergic pre-LC neurons exhibit spontaneous action potentials with different patterns, with a mean rate of up to 2 Hz (Kuo et al., 2023). One possible explanation is that the PRS (2 Hz) was unable to further affect the firing pattern of postsynaptic neurons that were already firing at a similar rate. Additionally, as previously discussed, the LC and various cortical regions have extensive mutual connections. Thus, GABAergic preLC neurons are not easily activated by a single cortical input but rather by integrating functionally correlated inputs. Together, these facts may explain discrepancies in previous *in vivo* studies. Some studies have found that mPFC input inhibits LC-NA neurons by recruiting local GABAergic neurons (Hervé-Minvielle & Sara, 1995; Breton-Provencher & Sur, 2019), while others have found no observable effect of inhibitory circuitry blockage on mPFC-to-NA neuron transmission in the LC (Jodo & Aston-Jones, 1997; Jodo et al., 1998). Due to methodological limitations, it should be kept in mind that the effect of mPFC input on LC-NA neurons and GABAergic preLC neurons in living animals may be much more significant than reported here. First, to increase the success rate of whole-cell recordings, we performed the recordings at room temperature (25°C) rather than at a more physiological temperature. This obviously reduces the impact of mPFC stimulation on LC-NA and GABAergic pre-LC neuron firing activity. Other factors, which are unavoidable for *ex vivo* recordings, worsen the situation. These include the much larger extracellular space *ex vivo*, which promotes the spread of electrical signals to the environment and results in the loss of neuronal electrical signals. Additionally, there are fewer healthy neuronal and glial cells ex vivo.

Despite these limitations, this study revealed important and genuine mechanisms. For instance, the mPFC exhibits monosynaptic connections with both NA and GABAergic preLC neurons. These inputs exhibit cell-type-specific differences in glutamate release properties. Compared to GABAergic preLC neurons, mPFC-to-LC-NA has a lower release probability (higher paired-pulse ratio), which enables presynaptic enhancement of glutamate release efficacy during behavior. The simultaneous connections of principal (LC-NA) and local (GABAergic preLC) neurons, which exhibit cell-type-specific differences in plastic function, enable the mPFC to effectively multiplex information to the LC for adaptive behavioral regulation.

## Acknowledgments

We are very grateful to Professor Y. Yanagawa for gifting the GAD^GFP^ mice. We also thank the technical support from the Technology Commons, College of Life Science, National Taiwan University. This work was supported by the National Science and and Technology Council, Taiwan [Grant Numbers: 111-2320-B-002-015-MY3 (MMY) and 110-2320-B-040-002-MY3 (HWY)].

## Author Contributions

PHL and WCH conducted the experiments, collected and analyzed the data and wrote the manuscript. HWY and MYM conceptualized the study, designed the experiments and wrote the manuscript.

## Data Availability Statement

All data supporting the findings of this study are available from the corresponding authors upon reasonable request.

## References

Arnsten, A.F.T. and Goldman-Rakic, P.S. (1984) Selective prefrontal cortical projections to the region of the locus coeruleus and raphe nuclei in the rhesus monkey. Brain Res. 306, 9–18.

Aston-Jones, G. and Bloom, F. E. (1981). Activity of norepinephrine-containing locus coeruleus neurons in behaving rats anticipates fluctuations in the sleep-waking cycle. J. Neurosci. 1, 876–86.

Aston-Jones, G., Rajkowski, J., Ivanova, S., Usher, M. and Cohen J. D. (1998). Neuromodulation and cognitive performance: recent studies of noradrenergic noradrenergic locus coeruleus neurons in behaving monkeys. Adv. Pharmacol. 42, 755–59.

Aston-Jones, G., Rajkowski, J. and Kubiak, P. (1997). Conditioned responses of monkey locus coeruleus neurons anticipate acquisition of discriminative behavior in a vigilance task. Neuroscience 80, 697–705.

Aston-Jones, G. and Waterhouse, B. D. (2016). Locus coeruleus: from global projection system to adaptive regulation of behavior. Brain Res. 1465, 75–78.

Aston-Jones, G., Zhu, Y. and Card, J. P. (2004). Numerous GABAergic afferents to locus coeruleus in the pericerulear dendritic zone: possible interneuronal pool. J. Neurosci. 24, 2313–21.

Berridge, C. W., Page, M. E., Valentino, R. J. and Foote, S. L. (1993). Effects of locus coeruleus inactivation on electroencephalographic activity in neocortex and hippocampus. Neuroscience 55, 381–93.

Berridge, C. W., Schmeichel, B. E. and España, R. A. (2018). Noradrenergic modulation of wakefulness/ arousal. Sleep Med. Rev. 16, 187–97.

Bouret S and Richmond BJ (2015) Sensitivity of Locus Ceruleus Neurons to Reward Value for Goal-Directed Actions. J Neurosci 35(9):4005–14.

Bouret, S. and Sara, S. J. (2004). Reward expectation, orientation of attention and locus coeruleus-mediated frontal cortex interplay during learning. Eur. J. Neurosci. 20, 791–802.

Bouret, S. and Sara, S. J. (2005). Network reset: a simplified overarching theory of locus coeruleus noradrenaline function. Trends Neurosci. 28, 574–82.

Breton-Provencher V and Sur M (2019) Active control of arousal by a locus coeruleus GABAergic circuit. Nat Neurosci 22:218–28.

Burdakov D, Ashcroft FM (2002) Cholecystokinin tunes firing of an electrically distinct subset of arcuate nucleus neurons by activating A-type potassium channels. J Neurosci 22:6380–6387.

Burdakov D, Alexopoulos H, Vincent A, Ashcroft FM (2004) Lowvoltage-activated A-current controls the firing dynamics of mouse hypothalamic orexin neurons. Eur J Neurosci 20:3281–3285.

Carter, M. E., Yizhar, O., Chikahisa, S., Nguyen, H., Adamantidis, A., Nishino, S., Deisseroth, K. and de Lecea, L. (2010). Tuning arousal with optogenetic modulation of locus coeruleus neurons. Nat. Neurosci. 13, 1526–33.

Clayton, E. C., Rajkowski, J., Cohen, J. D. and Aston-Jones, G. (2004). Phasic activation of monkey locus coeruleus neurons by simple decision in a forced-choice task. J. Neurosci. 24, 9914–20.

de Oliveira RB, Howlett MC, Gravina FS, et al. (2010) Pacemaker currents in mouse locus coeruleus neurons. Neuroscience, 170:166–77.

Grimm C, Duss SN, Privitera M, Munn BR, Karalis N, Frässle S, Wilhelm M, Patriarchi T, Razansky D, Wenderoth N, Shine JM, Bohacek J, Zerbi V. Tonic and burst-like locus coeruleus stimulation distinctly shift network activity across the cortical hierarchy. Nat Neurosci. 2024 Nov;27(11):2167–2177. doi: 10.1038/s41593-024-01755-8. Epub 2024 Sep 16. PMID: 39284964; PMCID: PMC11537968.

Hobson, J. A., McCarley, R. W. and Wyzinski, P. W. (1975). Sleep cycle oscillation: reciprocal discharge by two brainstem neuronal groups. Science 189, 55–58.

Jin, X., Li, S., Bondy, B., Zhong, W., Oginsky, M. F., Wu, Y., Johnson, C.M., Zhang, S., Cui, N. and Jiang, C. (2016). Identification of a group of GABAergic neurons in the dorsomedial area of the locus coeruleus. PLoS One 11, e0146470.

Jodo E and Aston-Jones G (1997) Activation of locus coeruleus by prefrontal cortex is mediated by excitatory amino acid inputs. Brain Res 768(1-2):327–32.

Jodo E, Chiang C, and Aston-Jones G (1998) Potent excitatory influence of prefrontal cortex activity on noradrenergic locus coeruleus neurons. Neuroscience 83(1):63–79.

Hervé-Minvielle A and Sara SJ (1995) Rapid habituation of auditory responses of locus coeruleus cells in anaesthetized and awake rats. Neuroreport 6(10):1363–8.

Kuo CC, Chan H, Hung WC, Chen RF, Yang HW, Min MY (2023) Carbachol increases locus coeruleus activation by targeting noradrenergic neurons, inhibitory interneurons and inhibitory synaptic transmission. Eur J Neurosci. 2023 Jan;57(1):32–53. doi: 10.1111/ejn.15866. Epub 2022 Dec 2. PMID: 36382388.

Kuo CC, Hsieh JC, Tsai HC, Kuo YS, Yau HJ, Chen CC, Chen RF, Yang HW, and Min MY (2020) Inhibitory interneurons regulate phasic activity of noradrenergic neurons in the mouse locus coeruleus and functional implications. J Physiol 598(18):4003–4029.

Liss B, Franz O, Sewing S, Bruns R, Neuhoff H, Roeper J (2001) Tuning pacemaker frequency of individual dopaminergic neurons by Kv4.3L and KChip3.1 transcription. EMBO J 20:5715–5724.

Luskin AT, Li L, Fu X, Martin MM, Barcomb K, Girven KS, Blackburn T, Wells BA, Thai ST, Li EM, Rana AN, Simon RC, Sun L, Gao L, Murry AD, Golden SA, Stuber GD, Ford CP, Gu L, Bruchas MR. Heterogeneous pericoerulear neurons tune arousal and exploratory behaviours. Nature. 2025 Jul;643(8071):437–447. doi: 10.1038/s41586-025-08952-w. Epub 2025 May 7. PMID: 40335695; PMCID: PMC12240712.

Min MY, Wu YW, Shih PY, Lu HW, Wu Y, Hsu CL, Li MJ, Yang HW. Roles of A-type potassium currents in tuning spike frequency and integrating synaptic transmission in noradrenergic neurons of the A7 catecholamine cell group in rats. Neuroscience. 2010 Jul 14;168(3):633–45. doi: 10.1016/j.neuroscience.2010.03.063. Epub 2010 Apr 8. PMID: 20381592.

Petreanu L, Mao T, Sternson S, Svoboda K. (2009) The subcellular organization of neocortical excitatory connections. Nature 457, 1142–45.

Poe GR, Foote S, Eschenko O, Johansen JP, Bouret S, Aston-Jones G, Harley CW, Manahan-Vaughan D, Weinshenker D, Valentino R, Berridge C, Chandler DJ, Waterhouse B, Sara SJ. Locus coeruleus: a new look at the blue spot. Nat Rev Neurosci. 2020 Nov;21(11):644–659. doi: 10.1038/s41583-020-0360-9. Epub 2020 Sep 17. PMID: 32943779; PMCID: PMC8991985.

Reyes A, Lujan R, Rozov A, Burnashev N, Somogyi P, Sakmann B. Target-cell-specific facilitation and depression in neocortical circuits. Nat Neurosci. 1998 Aug;1(4):279–85. doi: 10.1038/1092. PMID: 10195160.

Sesack SR, Deutch AY, Roth TH, and Bunney BS (1989) Topographical organization of the efferent projections of the medial prefrontal cortex in the rat: an anterograde tract-tracing study with Phaseolus vulgaris leucoagglutinin. J Comp Neurol 290(2):213–42.

Scanziani M, Gähwiler BH, Charpak S. Target cell-specific modulation of transmitter release at terminals from a single axon. Proc Natl Acad Sci U S A. 1998 Sep 29;95(20):12004–9. doi: 10.1073/pnas.95.20.12004. PMID: 9751780; PMCID: PMC21755.

Schwarz, L. A., Miyamichi, K., Gao, X. J., Bier, K. V., Weissbourd, B. D., DeLoach, K. E., Ren, J., Ibanes, S., Malenka, R. C., Kremer, E. J. and Luo, L. (2015). Viral-genetic tracing of the input-output organization of a central noradrenaline circuit. Nature 524, 88–92.

Swanson, L. W. and Hartman, B. K. (1975). The central adrenergic system. An immunofluorescence study of the location of cell bodies and their efferent connections in the rat utilizing dopamine-beta-hydroxylase as a marker. J. Comp. Neurol. 163, 467–505.

Takahashi K, Kayama Y, Lin JS, et al. (2010) Locus coeruleus neuronal activity during the sleep-waking cycle in mice. Neuroscience, 169, 1115–26.

Tóth K, and McBain CJ. (2000) Target-specific expression of pre-and postsynaptic mechanisms. J Physiol. 525, 41–51.

Tsunekawa N, Yanagawa Y & Obata K (2005). Development of GABAergic neurons from the ventricular zone in the superior colliculus of the mouse. Neurosci Res 51, 243–251.

Vazey EM, Moorman DE, and Aston-Jones G (2018) Phasic locus coeruleus activity regulates cortical encoding of salience information. PNAS 115, E9439–48.

Wang HY, Kuo ZC, Fu YS, Chen RF, Min MY and Yang HW (2015) GABAB receptor-mediated tonic inhibition regulates the spontaneous firing of locus coeruleus neurons in developing rats and in citalopram-treated rats. J Physiol 593:161–180.

Watanabe M, Uematsu A, and Johanses JP (2025) Bidirectional emotional regulation through prefrontal innervation of the locus coeruleus. Mol. Psychiatry 30, 3568–78.

Wilmot JH, Diniz CRAF, Crestani AP, Puhger KR, Roshgadol J, Tian L, and Wiltgen BJ (2024) Phasic locus coeruleus activity enhances trace fear conditioning by increasing dopamine release in the hippocampus. Elife 12:RP91465.

Wolfart J, Debay D, Masson GL, Destexhe A, and Bal T (2005) Synaptic background activity controls spike transfer from thalamus to cortex. Nat Neurosci 8(12):1760–7.

Zingg B, Chou XL, Zhang ZG, Mesik L, Liang F, Tao HW, and Zhang LI (2017) AAV-mediated Anterograde Transsynaptic Tagging: Mapping Input-Defined Functional Neural Pathways for Defense Behavior. Neuron 93:33–47.

